# Core Antibiotic-Induced Transcriptional Signatures Reflect Susceptibility to All Members of an Antibiotic Class

**DOI:** 10.1101/2020.10.29.359208

**Authors:** Melanie A. Martinsen, Alexis Jaramillo Cartagena, Roby P. Bhattacharyya

## Abstract

Current growth-based antibiotic susceptibility testing (AST) is too slow to guide early therapy. We previously developed a diagnostic approach that quantifies antibiotic-induced transcriptional signatures to distinguish susceptible from resistant isolates, providing phenotypic AST 24-36h faster than current methods. Here, we show that 10 transcripts optimized for AST of one fluoroquinolone, aminoglycoside, or beta-lactam reflect susceptibility to other members of that class. This finding will streamline development and implementation of this strategy, facilitating efficient antibiotic deployment.

---

Worldwide, more than 700,000 people die annually from antibiotic-resistant infections (1), and gaps in global antibiotic resistance tracking suggests this burden is severely underestimated (2, 3). Antibiotic-resistant infections lead to higher medical costs, longer hospital stays, and increased mortality (4, 5). Current growth-based antibiotic susceptibility testing (AST) is too slow to guide therapy in real time (6–8). This diagnostic delay causes overreliance on empiric broad-spectrum antimicrobials, contributing to the emergence of resistance and poor patient outcomes (9, 10). We recently developed a novel microbial diagnostic assay called GoPhAST-R (combined **G**en**o**typic and **Ph**enotypic **AST** through **R**NA detection) that can provide AST in <4 hours directly from blood culture, 24-36 hours faster than standard growth-based methods (11). GoPhAST-R quantifies specific antibiotic-responsive mRNA expression signatures using the commercially available RNA-detection platform, NanoString (12). After brief antibiotic exposure, susceptible cells become stressed, eliciting transcriptional changes that distinguish them from unharmed resistant cells. In addition to detecting these antibiotic-induced transcripts for phenotypic AST, GoPhAST-R can simultaneously target known genetic resistance markers to improve accuracy of resistance detection and facilitate molecular epidemiology.

We previously demonstrated GoPhAST-R for three antibiotics representing different classes, in five pathogens, and identified the top 10 antibiotic-responsive genes for each pathogen-antibiotic pair (11). To enhance flexibility of both development and implementation of this diagnostic, we aimed to test the generalizability of these transcriptional signatures of susceptibility within an antibiotic class. Since antibiotics elicit transcriptional responses related to their mechanism of action (13–16), we hypothesized that antibiotic-responsive genes optimized for one antibiotic could reflect susceptibility to all members of the same drug class. Here we confirm this hypothesis for two common pathogens with propensity for multidrug resistance – *Escherichia coli* and *Klebsiella pneumoniae* – treated with multiple members of three major antibiotic classes in regular clinical use – fluoroquinolones, aminoglycosides, and beta-lactams.

We targeted the top 10 genes we previously identified (11) for each of three individual antibiotics – ciprofloxacin (a fluoroquinolone, FQ), gentamicin (an aminoglycoside, AG), and meropenem (a beta-lactam, BL) – and assessed whether they reflect susceptibility to other members of their respective class (Table 1). All strains were obtained from clinical or reference microbiological laboratories, representing diverse geographic locations and resistance mechanisms when possible, and all reported minimum inhibitory concentrations (MICs) were verified by broth microdilution (6) (Dataset S1). For the FQ and AG classes, strains were grown in Mueller-Hinton Broth (MHB) and treated at early log phase. Some BLs exhibit inoculum effects, phenomena where the measured MIC depends on the number of microorganisms used in an AST assay (17–19); we previously showed that strains with reduced susceptibility due to inoculum effects also exhibit decreased induction of transcriptional susceptibility signatures at high cell density (11). To account for this effect, we grew all strains in MHB to early log phase, then diluted them to 2×10^5^ colony-forming units (cfu) ml^−1^ in fresh MHB and incubated for 60 minutes prior to treatment in order to achieve the CLSI-recommended MIC inoculum range of 2-8×10^5^ cfu ml^−1^. After brief antibiotic exposure (60 minutes for FQs and AGs, 120 minutes for BLs) at their respective clinical susceptibility breakpoint concentrations (6, 11, 20), or an equivalent control incubation without antibiotics, samples were mechanically lysed and used as input for NanoString assays as previously described (11). Using the NanoString platform (12), we quantified gene expression of both the top 10 responsive genes and 10 control genes we previously identified for individual pathogen-antibiotic pairs in multiplexed fashion (11). Control genes, whose expression is unaffected by antibiotics, were used to scale for cell number at lysis, and normalized fold-induction of each responsive gene in antibiotic-treated versus untreated samples was calculated for each strain, as previously described (11).

**TABLE 1.** Classification of antibiotics used in this study with respective treatment conditions

| Antibiotic | Class | Sub-class | Antibiotic treatment concentration<br>* (mg/L) | Antibiotic treatment duration<br>(minutes) |
| --- | --- | --- | --- | --- |
| ciprofloxacin | FQ | 2nd gen FQ | 0.25 | 60 |
| levofloxacin | FQ | 3rd gen FQ | 0.5 | 60 |
| moxifloxacin | FQ | 3rd gen FQ | 0.25 | 60 |
| gentamicin | AG |  | 4 | 60 |
| tobramycin | AG |  | 4 | 60 |
| amikacin | AG |  | 16 | 60 |
| ampicillin | BL | aminopenicillin | 8 | 120 |
| cefazolin | BL | 1st gen cephalosporin | 2 | 120 |
| ceftriaxone | BL | 3rd gen cephalosporin | 1 | 120 |
| aztreonam | BL | monobactam | 4 | 120 |
| piperacillin/tazobactam | BL | ureidopenicillin/ beta-lactamase inhibitor | 16/4 | 120 |
| ertapenem | BL | carbapenem | 0.5 | 120 |
| meropenem | BL | carbapenem | 1 | 120 |

\*The concentration for antibiotic treatment was chosen as the clinical susceptibility breakpoint established by the Clinical and Laboratory Standards Institute (CLSI) for all antibiotics whenever available (20), or the European Committee on Antimicrobial Susceptibility Testing (EUCAST) breakpoint for moxifloxacin (22).

We tested the ciprofloxacin 10-transcript signatures across the FQs ciprofloxacin, levofloxacin, and moxifloxacin, and the gentamicin 10-transcript signatures across the AGs gentamicin, tobramycin, and amikacin. For each class, we selected six isolates of each species: three susceptible and three resistant to all class members (Dataset S1). In both species, heatmaps illustrate that the top 10 genes identified for AST of ciprofloxacin and gentamicin showed similar normalized fold-induction upon treatment with three FQs and three AGs, respectively (Fig. 1a,b and Fig. S1a,b, top panels). One-dimensional projections summarizing these transcriptional data (11, 21) show robust distinction of susceptible and resistant isolates across each class (Fig. 1a,b and Fig. S1a,b, bottom panels).

**FIG 1.**
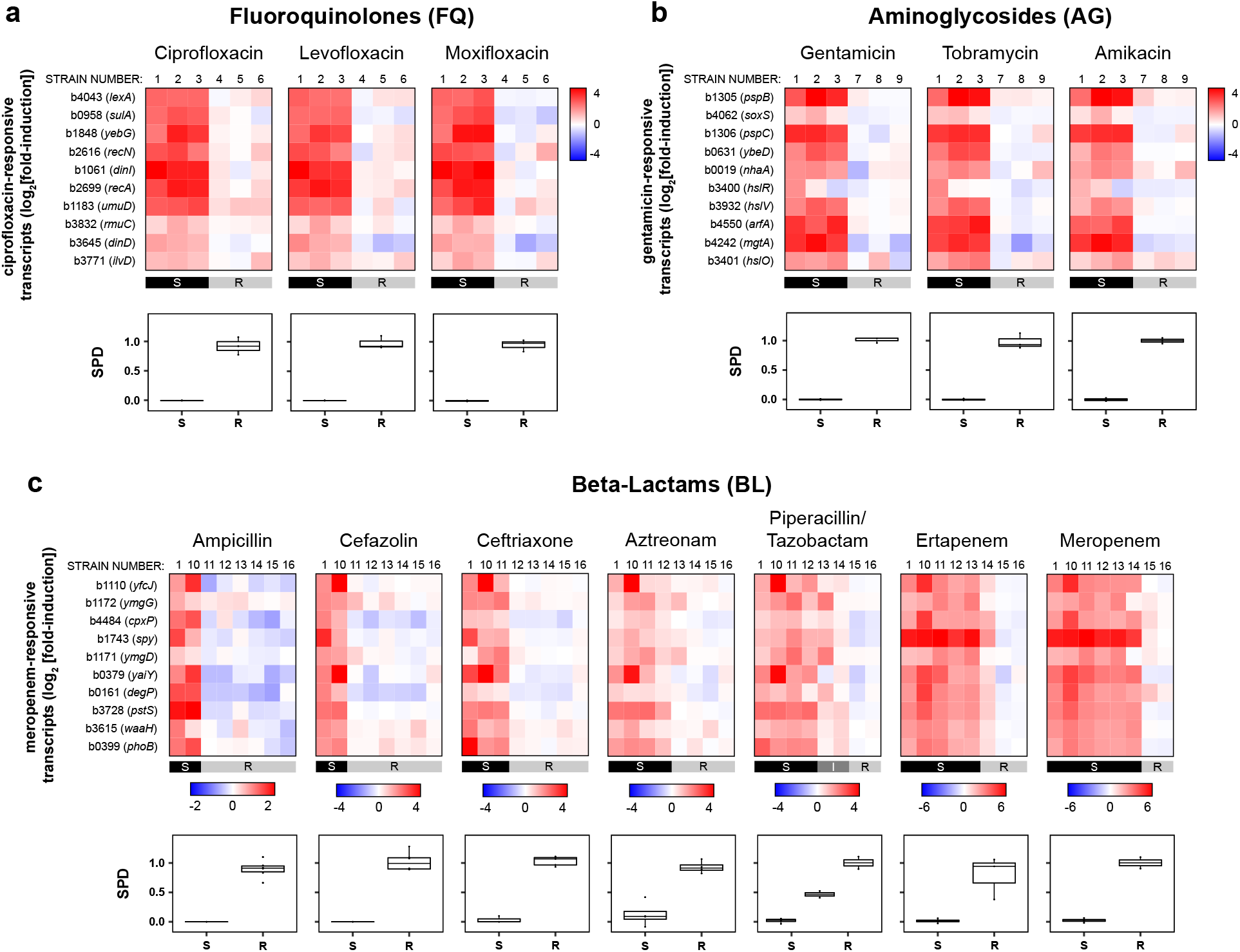
Differential expression of the same antibiotic-induced 10-transcript signatures from *E. coli* reflect susceptibility for all drugs within a class. Top panels show heatmaps of normalized, log-transformed fold-induction of the top 10 antibiotic-responsive transcripts we previously identified for *E. coli* treated with (**a**) ciprofloxacin, (**b**) gentamicin, and (**c**) meropenem upon exposure at CLSI breakpoint concentrations to other fluoroquinolones, aminoglycosides, and beta-lactams, respectively. Color-keys indicate range of log_2_[fold-induction] for transcripts in the respective heatmap(s). Strains are denoted by numbers over the heatmap columns, with CLSI classifications of each strain based on broth microdilution are shown below (S, susceptible; R, resistant). Gene identifiers for antibiotic-responsive transcripts are listed at left, as defined for NCBI reference sequence NC_000913. Bottom panels show one-dimensional projections (squared projected distance, SPD (11, 21)) of heatmap data for each strain, binned by CLSI classifications. By definition, an SPD of 0 indicates a transcriptional response to antibiotic equivalent to that of an average susceptible strain, while an SPD of 1 indicates a response equivalent to that of an average resistant strain. Data are summarized as box- and-whisker plots, where boxes extend from the 25^th^ to 75^th^ percentile for each category, with a line at the median, and whiskers extend from the minimum to the maximum.

We next tested BLs, a large class of diverse compounds comprising multiple subclasses that span a wide spectrum of anti-bacterial activity compared to FQs or AGs. The diversity of the BLs challenges the generalizability of transcriptional signatures of susceptibility within a drug class, and offers the most clinical benefit from rapid and efficient antibiotic deployment if successful. We tested the 10-transcript signatures identified for meropenem across treatments of seven BL antibiotics spanning multiple subclasses and ranging in spectrum of activity: ampicillin, cefazolin, ceftriaxone, aztreonam, piperacillin/tazobactam, ertapenem, and meropenem. We selected eight clinical isolates for each species that vary in susceptibility across the different BLs, ranging from pan-susceptible to pan-resistant (Dataset S1). Normalized fold-induction of the meropenem-responsive genes tested across the seven BLs are shown as heatmaps and summarized as one-dimensional projections (Fig. 1c, Fig. S1c). In both species, each BL induces the top 10 meropenem-responsive transcripts in only the susceptible isolates, allowing susceptibility distinction despite variability in expression levels of certain genes across the class. Strains with MICs closer to the breakpoint exhibit partial induction, consistent with our previous finding that the magnitude of transcriptional response to antibiotic exposure at the breakpoint correlates with MIC (11). Notably, strain 14 is correctly distinguishable from susceptible isolates under ertapenem exposure, despite lacking genotypic markers of common carbapenemases and extended-spectrum beta-lactamases (Fig. S2). This exemplifies GoPhAST-R’s ability to determine phenotypic resistance, agnostic to resistance mechanism.

This work demonstrates that the same antibiotic-induced 10-transcript signatures reflect antibiotic susceptibility for all drugs within a class, consistent with a conserved core transcriptional response to each of three major antibiotic classes. Despite their diversity, even BL compounds share a susceptibility signature across all subclasses, implying a common stress response to their similar cellular targets. The 10-transcript signatures used in this study, previously designed for individual antibiotics, may not represent the top 10 genes for the whole class nor for each individual member, but they clearly report on susceptibility for each compound in a given class. Moreover, this study provides further evidence for transcriptional profiling as a robust phenotypic measure of antimicrobial stress, underscoring the flexibility and breadth of GoPhAST-R: the same minimal gene set derived for one specific antibiotic can assess susceptibility not only across diverse strains but also across drugs with a shared mechanism of action. This finding will streamline GoPhAST-R implementation, contributing to the critical effort to employ rapid AST diagnostics to guide upfront selection of the narrowest effective agent against a given pathogen. Efficient, informed deployment of antibiotics will improve patient outcomes while minimizing selection for resistance.

## Acknowledgements

This work was supported in part by the National Institute of Allergy and Infectious Diseases of the National Institutes of Health under award 1K08AI119157-04 (R.P.B.), and in part by the Massachusetts General Hospital’s Pilot Translational Research Grant. The funders had no role in study design, data collection and interpretation, or the decision to submit the work for publication. The content is solely the responsibility of the authors and does not necessarily represent the official views of the National Institutes of Health.

We thank Gowtham Thakku, James Gomez, Jonathan Livny, and Peijun Ma for helpful discussion, Lorrie He for laboratory support during data collection, and Ashlee Earl, the CDC ARBank, and the New York State Department of Health for strains.

M.A.M. contributed to the project conceptualization, data curation, formal analysis, investigation, methodology, validation, visualization, and writing (original draft, review, and editing). A.J.C. contributed to a portion of the investigation as well as writing (review and editing). R.P.B. contributed to project conceptualization, formal analysis, funding acquisition, methodology, supervision, validation, and writing (review and editing). All authors have read and approved the manuscript

## References

1. O’Neill J. 2016. Tackling drug-resistant infections globally: final report and recommendations. Review on Antimicrobial Resistance.

2. WHO. 2014. Antimicrobial resistance: global report on surveillance. Geneva: World Health Organization.

3. WHO. 2020. Global antimicrobial resistance surveillance system (GLASS) report: early implementation 2020., Geneva: World Health Organization. Licence: CC BY-NC-SA 3.0 IGO.

4. Mauldin PD, Salgado CD, Hansen IS, Durup DT, Bosso JA. 2010. Attributable Hospital Cost and Length of Stay Associated with Health Care-Associated Infections Caused by Antibiotic-Resistant Gram-Negative Bacteria. Antimicrob Agents Chemother 54:109–115.

5. Kadri SS, Adjemian J, Lai YL, Spaulding AB, Ricotta E, Prevots DR, Palmore TN, Rhee C, Klompas M, Dekker JP, Powers JH3rd,, Suffredini AF, Hooper DC, Fridkin S, Danner RL. 2018. Difficult-to-Treat Resistance in Gram-negative Bacteremia at 173 US Hospitals: Retrospective Cohort Analysis of Prevalence, Predictors, and Outcome of Resistance to All First-line Agents. Clin Infect Dis 67:1803–1814.

6. Wiegand I, Hilpert K, Hancock REW. 2008. Agar and broth dilution methods to determine the minimal inhibitory concentration (MIC) of antimicrobial substances. Nat Protoc 3:163–175.

7. Caliendo AM, Gilbert DN, Ginocchio CC, Hanson KE, May L, Quinn TC, Tenover FC, Alland D, Blaschke AJ, Bonomo RA, Carroll KC, Ferraro MJ, Hirschhorn LR, Joseph WP, Karchmer T, MacIntyre AT, Reller LB, Jackson AF. 2013. Better tests, better care: improved diagnostics for infectious diseases. Clin Infect Dis 57 Suppl 3:S139–70.

8. Burnham CD, Leeds J, Nordmann P, O’Grady J, Patel J. 2017. Diagnosing antimicrobial resistance. Nat Rev Microbiol 15:697–703.

9. Rhee C, Kadri SS, Dekker JP, Danner RL, Chen HC, Fram D, Zhang F, Wang R, Klompas M. 2020. Prevalence of Antibiotic-Resistant Pathogens in Culture-Proven Sepsis and Outcomes Associated With Inadequate and Broad-Spectrum Empiric Antibiotic Use. JAMA Netw Open 3:e202899.

10. Kadri SS, Lai YL, Warner S, Strich JR, Babiker A, Ricotta EE, Demirkale CY, Dekker JP, Palmore TN, Rhee C, Klompas M, Hooper DC, Powers JH3rd,, Srinivasan A, Danner RL, Adjemian J. 2020. Inappropriate empirical antibiotic therapy for bloodstream infections based on discordant in-vitro susceptibilities: a retrospective cohort analysis of prevalence, predictors, and mortality risk in US hospitals. Lancet Infect Dis doi:10.1016/s1473-3099(20)30477-1.

11. Bhattacharyya RP, Bandyopadhyay N, Ma P, Son SS, Liu J, He LL, Wu L, Khafizov R, Boykin R, Cerqueira GC, Pironti A, Rudy RF, Patel MM, Yang R, Skerry J, Nazarian E, Musser KA, Taylor J, Pierce VM, Earl AM, Cosimi LA, Shoresh N, Beechem J, Livny J, Hung DT. 2019. Simultaneous detection of genotype and phenotype enables rapid and accurate antibiotic susceptibility determination. Nat Med 25:1858–1864.

12. Geiss GK, Bumgarner RE, Birditt B, Dahl T, Dowidar N, Dunaway DL, Fell HP, Ferree S, George RD, Grogan T, James JJ, Maysuria M, Mitton JD, Oliveri P, Osborn JL, Peng T, Ratcliffe AL, Webster PJ, Davidson EH, Hood L, Dimitrov K. 2008. Direct multiplexed measurement of gene expression with color-coded probe pairs. Nat Biotechnol 26:317–25.

13. O’Rourke A, Beyhan S, Choi Y, Morales P, Chan AP, Espinoza JL, Dupont CL, Meyer KJ, Spoering A, Lewis K, Nierman WC, Nelson KE. 2020. Mechanism-of-Action Classification of Antibiotics by Global Transcriptome Profiling. Antimicrob Agents Chemother 64:e01207–19.

14. Boshoff HIM, Myers TG, Copp BR, McNeil MR, Wilson MA, Barry 3rd CE. 2004. The Transcriptional Responses of Mycobacterium tuberculosis to Inhibitors of Metabolism: Novel Insights into Drug Mechanisms of Action. J Biol Chem doi:10.1074/jbc.M406796200:40174-40184.

15. Brazas MD, Hancock RE. 2005. Using microarray gene signatures to elucidate mechanisms of antibiotic action and resistance. Drug Discov Today 10:1245–52.

16. Hutter B, Schaab C, Albrecht S, Borgmann M, Brunner NA, Freiberg C, Ziegelbauer K, Rock CO, Ivanov I, Loferer H. 2004. Prediction of Mechanisms of Action of Antibacterial Compounds by Gene Expression Profiling. Antimicrob Agents Chemother 48:2838–2844.

17. Smith KP, Kirby JE. 2018. The Inoculum Effect in the Era of Multidrug Resistance: Minor Differences in Inoculum Have Dramatic Effect on MIC Determination. Antimicrob Agents Chemother 62:e00433–18.

18. Eng RH, Cherubin C, Smith SM, Buccini F. 1985. Inoculum effect of beta-lactam antibiotics on Enterobacteriaceae. Antimicrob Agents Chemother 28:601–606.

19. Brook I. 1989. Inoculum Effect. Rev Infect Dis 11:361–368.

20. CLSI. 2020. Performance Standards for Antimicrobial Susceptibility Testing. Wayne, PA: Clinical and Laboratory Standards Institute.

21. Barczak AK, Gomez JE, Kaufmann BB, Hinson ER, Cosimi L, Borowsky ML, Onderdonk AB, Stanley SA, Kaur D, Bryant KF, Knipe DM, Sloutsky A, Hung DT. 2012. RNA signatures allow rapid identification of pathogens and antibiotic susceptibilities. Proc Natl Acad Sci U S A 109:6217–6222.

22. EUCAST. 2020. Breakpoint tables for interpretation of MICs and zone diameters. Version 10.0. The European Committee on Antimicrobial Susceptibility Testing.

