## Supplemental Figures S1 and S2 for "Core Antibiotic-Induced Transcriptional Signatures Reflect Susceptibility to All Members of an Antibiotic Class"

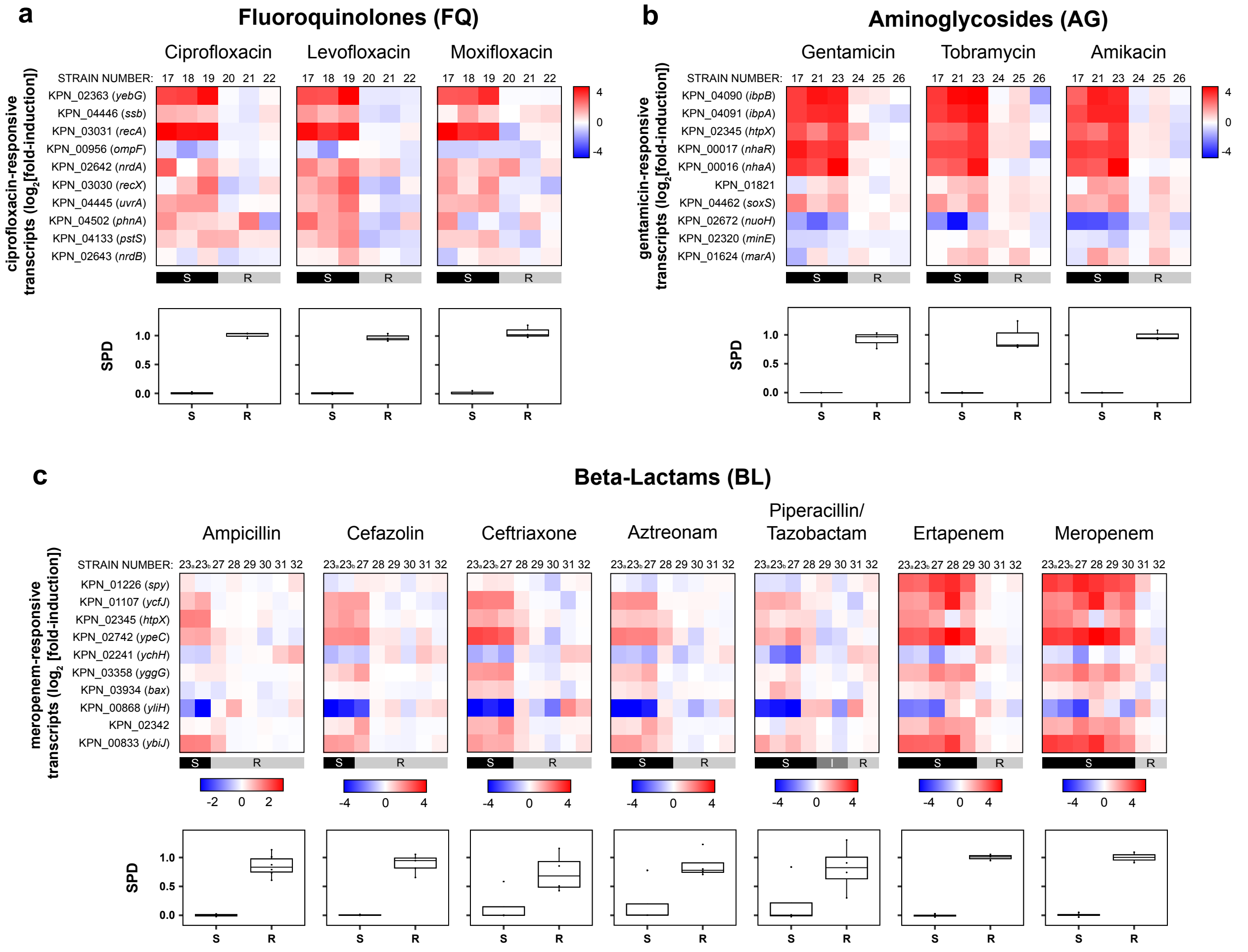


**FIG S1. Differential expression of the same antibiotic-induced 10-transcript signatures from *K. pneumoniae* reflect susceptibility for all drugs within a class.** Top panels show heatmaps of normalized, log-transformed fold-induction of the top 10 antibiotic-responsive transcripts we previously identified for *K. pneumoniae* treated with (**a**) ciprofloxacin, (**b**) gentamicin, and (**c**) meropenem upon exposure at CLSI breakpoint concentrations to other fluoroquinolones, aminoglycosides, and beta-lactams, respectively. Color-keys indicate range of $log_2$[fold-induction] for transcripts in the respective heatmap(s). Strains are denoted by numbers over the heatmap columns, with CLSI classifications of each strain based on broth microdilution are shown below (S, susceptible; R, resistant). Gene identifiers for antibiotic-responsive transcripts are listed at left, as defined for NCBI reference sequence NC_009648. Bottom panels show one-dimensional projections (squared projected distance, SPD (1, 2)) of heatmap data for each strain, binned by CLSI classifications. By definition, an SPD of 0 indicates a transcriptional response to antibiotic equivalent to that of an average susceptible strain, while an SPD of 1 indicates a response equivalent to that of an average resistant strain. Data are summarized as box-and-whisker plots, where boxes extend from the 25$^{th}$ to 75$^{th}$ percentile for each category, with a line at the median, and whiskers extend from the minimum to the maximum.

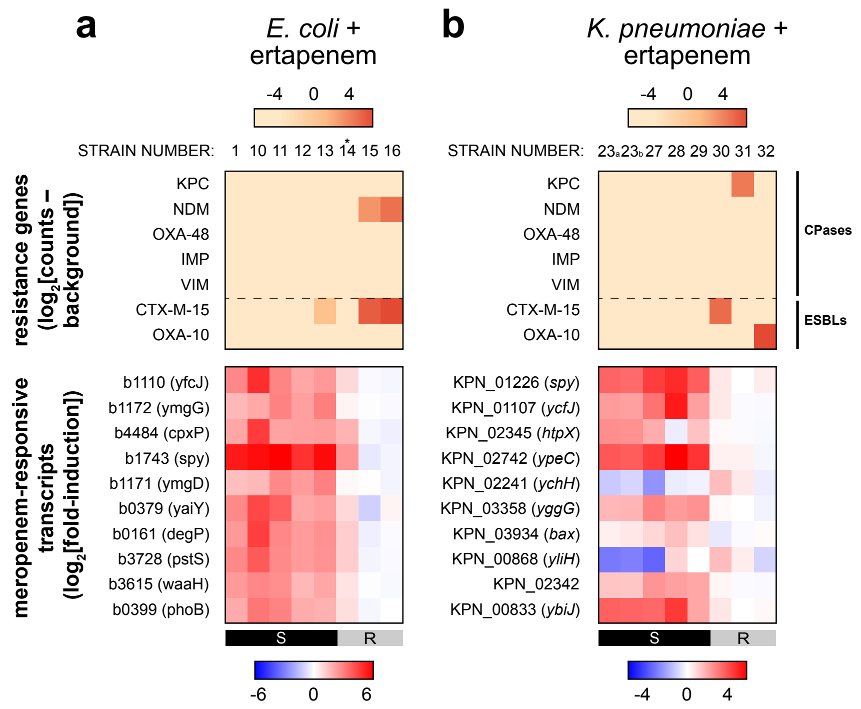


**FIG S2. GoPhAST-R detects carbapenemase (CPase) and extended-spectrum beta-lactamase (ESBL) gene content, augmenting phenotypic AST.** Top panels show GoPhAST-R detection of CPase and select ESBL transcript content in (**a**) *E. coli* and (**b**) *K. pneumoniae* strains selected for BL treatment. Heatmap intensity reflects normalized, background-subtracted, log-transformed NanoString data from probes for the indicated gene families, as described (1). Color-keys indicate range of $\log_2$[counts – background] for these genes in the respective heatmap(s). Bottom panels show the GoPhAST-R phenotypic AST heatmaps for ertapenem treatment from (**a**) Fig. 1c and (**b**) Fig. S1c. *Note strain 14 displays phenotypic ertapenem resistance without detectable CPase or ESBL.

### References

1. Bhattacharyya RP, Bandyopadhyay N, Ma P, Son SS, Liu J, He LL, Wu L, Khafizov R, Boykin R, Cerqueira GC, Pironti A, Rudy RF, Patel MM, Yang R, Skerry J, Nazarian E, Musser KA, Taylor J, Pierce VM, Earl AM, Cosimi LA, Shoresh N, Beechem J, Livny J, Hung DT. 2019. Simultaneous detection of genotype and phenotype enables rapid and accurate antibiotic susceptibility determination. Nat Med 25:1858-1864.
2. Barczak AK, Gomez JE, Kaufmann BB, Hinson ER, Cosimi L, Borowsky ML, Onderdonk AB, Stanley SA, Kaur D, Bryant KF, Knipe DM, Sloutsky A, Hung DT. 2012. RNA signatures allow rapid identification of pathogens and antibiotic susceptibilities. Proc Natl Acad Sci U S A 109:6217-6222.
